# The importance of neutral over niche processes in structuring Ediacaran early animal communities

**DOI:** 10.1101/443275

**Authors:** Emily G. Mitchell, Simon Harris, Charlotte G. Kenchington, Philip Vixseboxse, Lucy Roberts, Catherine Clark, Alexandra Dennis, Alexander G. Liu, Philip R. Wilby

## Abstract

The relative influence of niche versus neutral processes in ecosystem dynamics is a fundamental question in community ecology, but the extent to which they structured early animal communities is unknown. The oldest known metazoan-dominated paleocommunities occur in Ediacaran age (~565 million years old) strata in Newfoundland, Canada and Charnwood Forest, UK. These comprise large and diverse in-situ populations of sessile organisms that are amenable to spatial point process analyses, enabling inference of the most likely underlying niche or neutral processes governing their community structure. We conducted comprehensive spatial mapping of seven of the largest Ediacaran paleocommunities using LiDAR, photogrammetry and a laser-line probe. We find neutral processes to dominate these paleocommunities with limited influence of niche processes. Our results differ from the niche-dominated dynamics of modern marine ecosystems, revealing that the dynamics of environmental interactions prompted very different ecosystem structuring for these early animal communities.

## Introduction

Two opposing theories lie at the heart of debate regarding the fundamental mechanisms that govern ecosystem structure and biodiversity: niche and neutral. Niche theory is a central tenet of classical ecological theory, whereby species avoid competitive exclusion by occupying different niches within the ecosystem (1). Smaller niche overlaps result in less competition between taxa, permitting numerous taxa to exist in an area without excluding each other. Species are able to co-exist because they have different requirements. Niche models describe selection-dominated ecosystems, whereby species dynamics operate deterministically as a series of inter-specific interactions, which act as stabilizing mechanisms for the ecosystem (2).

Neutral processes are often referred to as the ‘null model’ of niche processes: instead of species *differences* enabling co-existence, it is their *similarities* that drive high diversity (3). Within neutral models, species fitness is similar across a community, and so different taxa can co-exist because no single taxon has a significant competitive advantage over any other. Despite this seemingly unrealistic assumption, neutral theories have been able to accurately reproduce certain species-area-distributions (3) and beta diversity patterns (4, 5), sometimes better than niche theories (1).

In recent years, unified or continuous theories have emerged, whereby niche and neutral processes combine to enable species coexistence (2, 6). In these combined models, species can exhibit strong differences and strong stabilizations (niche-type), or similar fitness and weak stabilizations (neutral-type), with the classic niche and neutral models forming the extreme end-members of this continuum model. However, it is often not possible to select the best-fit niche or neutral model, making it difficult to disentangle the relative influence of niche-and neutral-type processes within modern, complex ecosystems (6).

In order to investigate how niche and neutral processes contributed to community dynamics in deep time, we focus on some of the oldest macroscopic metazoan-dominated paleocommunities that are currently known: those comprising the Avalonian Assemblage of the Ediacaran macrobiota (7 – 9). The evolution of macroscopic metazoans was coupled with a transformation in ecosystem dynamics, with paleocommunities evolving from pre-Ediacaran microbial populations with assumed simple community structure (10), via late Ediacaran (571–540 Ma) paleocommunities that exhibited both simple and complex community structures (11), into Cambrian ‘modern’ metazoan ecosystems with comparable ecosystem structures to the present day (12). Some of the oldest metazoan-dominated communities form part of the Avalonian Assemblage of the Ediacaran macrobiota (13), and are known primarily from Newfoundland, Canada and Leicestershire, UK (Fig. S1). Avalonian soft-bodied organisms were sessile and preserved *in-situ* in deep-water paleoenvironments dated to ~571–560 Ma (14, 15), beneath volcanogenic/volcaniclastic event beds (16, 17). As such, bedding-plane surfaces exposed by modern weathering of tuffs preserve near-complete census paleocommunities (16, 18), although the impact of erosion of these surfaces needs to be considered (19; cf. 20). Since they were soft bodied, dead organisms could not accumulate over long time periods, reducing the extent of time-averaging. Avalonian ecosystems pre-date macro-predation and vertical burrowing, and so remained in place post-mortem (21–23), with no evidence of locomotion on any of these bedding-planes. Consequently, the size and position of each specimen can be considered an accurate record of the organism’s life history, including its dispersal and the habitat and community interactions it was subject to. In common with living communities, it is therefore possible to use spatial point process analyses (SPPA) to infer the most likely underlying ecological and biological processes in operation (25).

For sessile organisms, community-scale spatial distributions depend on the interplay of a limited number of different factors, namely physical environment (which manifests as habitat associations of a taxon or taxon-pairs; 26), organism dispersal/reproduction (27), competition for resources (28), facilitation between taxa (29), and differential mortality (30).

To assess the relative influence of niche and neutral processes for sessile communities, niche processes are identified as intra-or inter-specific habitat associations, and/or intra-and inter-specific competition and/or facilitation (31). Neutral processes are identified where univariate distributions exhibit complete spatial randomness (CSR), and by dispersal processes that are independent of local environment (i.e. habitat heterogeneities; 31–35). Dispersal patterns are indicated by best-fit Thomas Cluster (TC) or Double Thomas Cluster (DTC) models (31).

CSR indicates neutral processes because there are no biologically or ecologically significant intrinsic or extrinsic influences on the spatial distribution. TC and DTC aggregations are also considered neutral since they describe dispersal processes, whereby aggregations arise from propagules only traveling a limited distance, thus being unable to reach all suitable substrates regardless of underlying habitat heterogeneities or species requirements (27, 36, 37).

Intra-specific habitat associations are best-modelled by a heterogeneous Poisson model (HP), or when combined with dispersal limitations, an Inhomogeneous Thomas Cluster model (ITC; 31, 37). Density-dependent competition, as indicated by size-dependent spatial segregation (38), indicates a lack of sufficient resources, and is therefore a niche-based process. The other bivariate or inter-specific interactions between taxa include facilitation, which is considered niche because the requirement of one taxon relying on another indicates that the facilitated taxon could not survive independently. Facilitation is best modelled by a Linked Cluster model (LCM). Habitat associations between taxa are also considered niche processes because such taxa associations correspond to the underlying habitat variations on which the species depend, and are best modelled by a shared parents models (SPM) and/or heterogeneous Poisson models (22). Therefore, for univariate distributions, neutral processes are indicated by CSR, TC or DTC models, and niche processes by segregation and HP and ITC models (Fig. S3). For bivariate distributions, neutral processes are indicated by CSR while niche processes are indicated by segregation, LCM, SPM and/or HP models.

## Methods

### Data collection and extraction

In this study we assessed the univariate and bivariate spatial distributions of taxa from seven Avalonian bedding-plane assemblages: the ‘D’, ‘E’, and Bristy Cove (X-Ray) surfaces in the Mistaken Point Ecological Reserve, the St. Shott’s surface at Sword Point; the H14 (Johnson) surface at Little Catalina and Spaniard’s Bay all in Newfoundland, Canada; and Bed B in Charnwood Forest, UK (Fig. S1, Table S1). These spatial analyses require the mapping of large spatial areas (up to 115m^2^), in sufficient resolution to be able to taxonomically identify the specimens from the resulting digital dataset. The best way to map the surfaces differed depending on the preservation and dip of the surface. All surfaces were LiDAR scanned using a Faro Focus 330X to ensure spatial accuracy was maintained over large areas. The LiDAR scans resulted in a 3D surface mesh of 1 mm resolution. The Spaniard’s Bay and Mistaken Point ‘D’ and ‘E’ surfaces were laser scanned using a Faro Scan Arm LLP, resulting in surface meshes of 0.050 mm resolution. The high-resolution scanning was done in grids of 1m × 1m. Due to large file sizes, these high-resolution scans could not all be viewed simultaneously, so control points were marked in both each high-resolution scan, and in the LiDAR scan, enabling accurate combination of the high-resolution scans with the LiDAR surface data (done using Geomagic 2015). Taxon identification, position, and fossil dimensions of disc width, disc length, stem length, stem width, frond length and frond width were marked up in Inkscape 0.92.3 on a 2D map of the combined dataset as vectors for every specimen, creating a 2D vector map of the paleocommunity.

For H14, Bristy Cove and St Shott’s surfaces, fossil relief was not sufficiently high to permit accurate capture of all morphological details using the laser-line probe. These surfaces did have good colour differentiation of the fossils, so a photomap was created by photographing the specimens along a horizontal and vertical grid, and using Agisoft Photoscan software v1.3.5 to create a photogrammetric render of the surface. The LiDAR scan was then imported into Photoscan, and the photographs aligned on the LiDAR scan to ensure large-scale accuracy. An orthomosaic of the surface was produced within Agisoft PhotoScan, and the fossils marked up as vectors as above.

Bed B, Charnwood Forest has a dip of 45°, and so is not suitable for *in situ* high-resolution scanning using our equipment. Instead, we used Reflectance Transformation Images (RTIs) of casts of this surface (57, 58). Each RTI was marked up as a vector map and imported onto the LiDAR scan. The LiDAR scan enabled checking of mould deformation, and where needed was used to retrodeform the vector map.

Upon completion, vector maps were processed using a custom script in Haskell (59), which output the specimen identification number, taxonomic identification, and specimen dimensions. This output formed the basic dataset for the spatial analyses.

### Taxonomic identification

Specimens were assigned to one of twenty macrofossil taxa/groups, including several ‘bin’ groups (60), on the basis of their morphological attributes: 1) *Arborea*, 2) *Aspidella*, 3) *Avalofractus*, 4) *Beothukis*, 5) *Bradgatia*, 6) Brushes, 7) *Charnia*, 8) *Charniodiscus*, 9) “Feather Dusters” which includes *Plumeropriscum* and *Primocandlebrum*, 10) *Fractofusus andersoni* + *F*. *misrai*, 11) *Hylaecullulus*, 12) Ivesheadiomorphs, 13) Ostrich Feather, 14) *Pectinifrons*, 15) *Primocandelabrum*, 16) *Thectardis*, 17) *Trepassia*, 18) *Vinlandia*, 19) “Holdfast Discs” [all discoidal specimens of uncertain affinity, with or without associated stems, which lack sufficient detail to identify the taxon], 20) “Other Species” [rare forms that do not fall into any of the other groups; e.g., *Hapsidophyllas*]. Non-abundant taxa (taken as < 30 specimens) and taphomorphs (organ taxa, such as *Hiemalora* or the decayed remains of already dead organisms, such as ivesheadiomorphs) were excluded from analyses, leaving 13 abundant taxa, three of which (*Charniodiscus*, *Charnia*, *Bradgatia*) occur abundantly on two bedding-planes and one (*Fractofusus*) on four bedding-planes.

### Bias analyses

Differential erosion has the potential to distort spatial analyses (17, 19) so for each surface, we tested for erosional biases (19) and tectonic deformation, corrected for these factors into account if they significantly affected specimen density distributions (Fig. S2, Table S2). ․ Our data have been tested for the influence of differential erosion using heterogeneous Poisson models. We modelled possible sources of erosion (cf. 20), fitting at least three heterogeneous Poisson models to the data, with the models dependent on *x* (parallel to strike), *y* (parallel to dip), and a point chosen on a surface-by-surface basis dependent on the most likely point of erosion. The St. Shott’s, Bed B and H14 surfaces all showed significant fossil density changes depending on these physical features of the surface, implying that the observed surface specimen density has been significantly influenced by post-preservational erosion processes. Such heterogeneous erosional processes were incorporated into subsequent analyses so that the underlying biological and ecological processes could be investigated. Tectonically distorted data were retrodeformed by returning elongated holdfast discs to a circular outline (16, 20).

### Spatial Analyses

Initial data exploration, inhomogeneous Poisson modelling and segregation tests were performed in R (61) using the package spatstat (62–64). Programita was used to find distance measures and to perform aggregation model fitting (described in detail in references (65–68)).

The univariate spatial distribution of each taxon on each bedding plane was described using pair correlation functions (PCFs). A PCF = 1 indicates a distribution that was completely spatially random (CSR); PCF > 1 indicates aggregation; and PCF < 1 indicates segregation (25, 32, 39). Univariate and Bivariate pair correlation functions (PCFs) were calculated from the population density using a grid of 10cm × 10cm cells on all surfaces except Bristy Cove, where a 1cm × 1cm cell size was used. To minimise noise, a smoothing was applied to the PCF dependent on specimen abundance: This smoothing was over three cells with all surfaces except Bristy Cove which had a 5 cell smoothing. To test whether the PCF exhibited complete spatial randomness (CSR), 999 simulations were run for each univariate and bivariate distribution, with the 49^th^ highest and lowest values removed (69). CSR was modelled by a Poisson model on a homogeneous background where the PCF = 1 and the fit of the fossil data to CSR was assessed using Diggle’s goodness-of-fit test (32, 39). Note that due to non-independence of spatial data, Monte-Carlo generated simulation envelopes cannot be interpreted as confidence intervals. If the observed data fell below the Monte-Carlo simulations, the bivariate distribution was interpreted to be segregated; above the Monte-Carlo simulations, the bivariate distribution was found to be aggregated.

If a taxon was not randomly distributed on a homogeneous background, and was aggregated (Fig. S3, Table S3), the random model on a heterogeneous background was tested by creating a heterogeneous background created from the density map of the taxon under consideration, being defined by a circle of radius R over which the density is averaged throughout the sample area. Density maps were formed using estimators within the range of 0.1m < R < 1m, with R corresponding to the best-fit model used. If excursions outside the simulation envelopes for both homogeneous and heterogeneous Poisson models remained, then Thomas cluster models were fitted to the data as follows:

1. The PCF and L function (70) of the observed data were found. Both measures were calculated to ensure that the best-fit model is not optimized towards only one distance measure, and thus encapsulates all spatial characteristics.
2. Best-fit Thomas cluster processes (71) were fitted to the two functions where PCF>1. The best-fit lines were not fitted to fluctuations around the random line of PCF=1 in order to aid good fit about the actual aggregations, and to limit fitting of the model about random fluctuations. Programita used the minimal contrast method (32, 39) to find the best-fit model.
3. If the model did not describe the observed data well, the lines were re-fitted using just the PCF. If that fit was also poor, then only the L-function was used.
4. 99 simulations of this model were generated to create simulation envelopes, and the fit checked using the O-ring statistic. (64)
5. *p*_*d*_ was calculated over the model range. Very small-scale segregations (under 2 cm) were not included in the model fitting, since they likely represent the finite size of the specimens, and a lack of specimen overlap.
6. If there were no excursions outside the simulation envelope and the *p*_*d*_-value was high, then a univariate homogeneous Thomas cluster model was interpreted as the best model. For each bivariate distribution displaying segregation, the size-classes of each taxon were calculated, the bivariate PCFs of the smallest size-classes and largest size-classes were plotted, with 999 Monte Carlo simulations of a complete spatially random distribution and segregation tests performed. The most objective way to resolve the number and range of size classes in a population is by fitting height-frequency distribution data to various models, followed by comparison of (logarithmically scaled) Bayesian information criterion (BIC) values (72), which we performed in R using the package MCLUST (73). The number of populations thus identified was then used to define the most appropriate size classes. A BIC value difference of >10 corresponds to a “decisive” rejection of the hypothesis that two models are the same, whereas values <6 indicate only weakly rejected similarity of the models (72–77). Once defined, the PCFs for each size class were calculated. Although it was necessary to set firm boundaries for each size class, the populations are normally distributed and therefore overlap. As a result, the largest individuals of the small population are grouped within the middle size class, while some of the smallest of the medium population are included within the small size class. As such, the medium population was excluded from analyses.

## Results

Across the seven surfaces and the 19 taxon univariate distributions examined, eight taxon distributions were best modelled by CSR (Figs. 1 and 2, Table S3). Of the non-CSR taxon distributions, 10 were best modelled by TC (or DTC). Only *Trepassia* on Spaniard’s Bay was best modelled by an ITC model (Fig. 2, Table S3). The only taxa which had univariate spatial distribution with a HP best-fit model was *Beothukis* on the ‘E’ and Spaniard’s Bay surfaces (Figs. 1 and 2).

**Figure 1.**
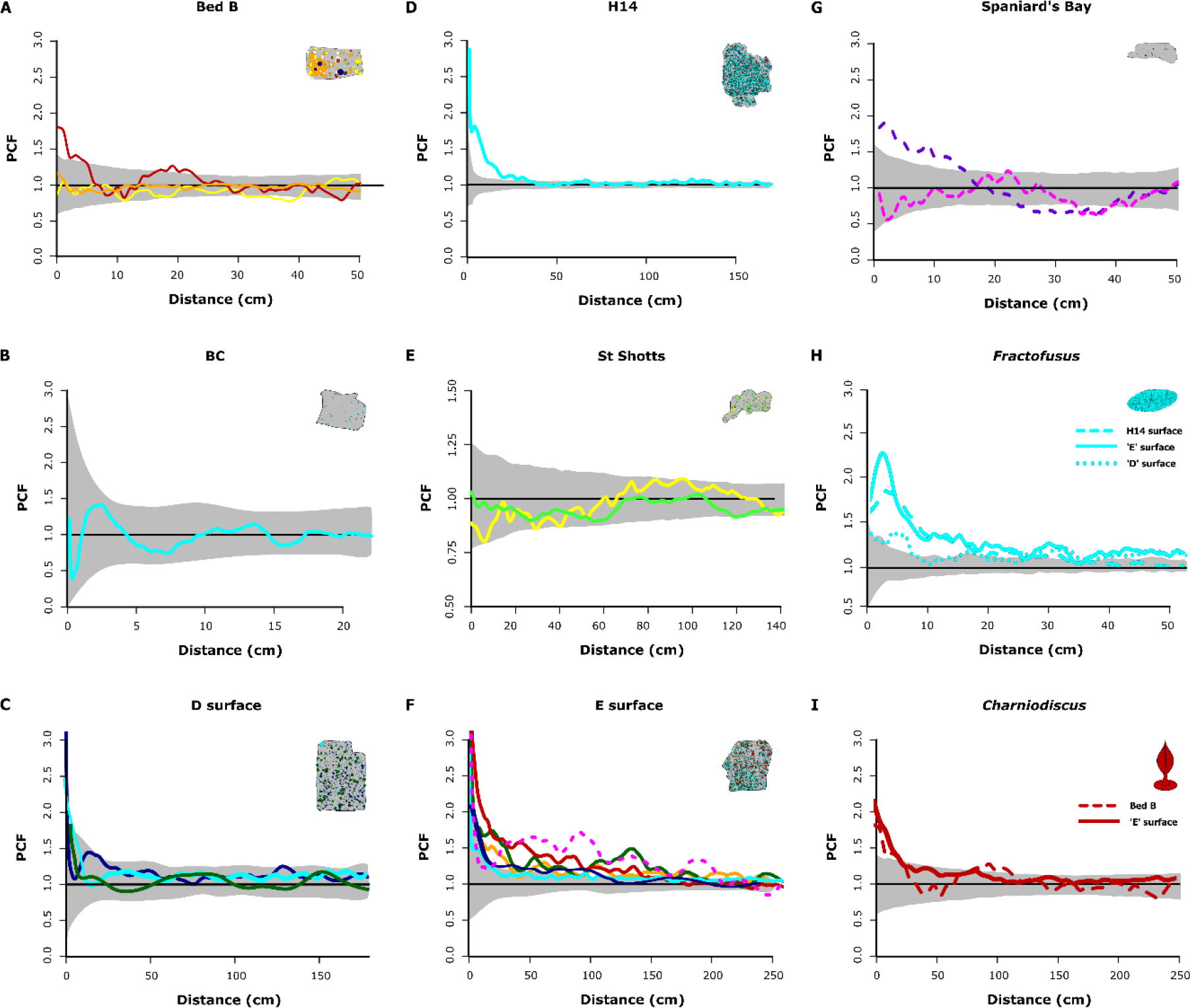
Univariate PCF for the seven Ediacaran fossil surfaces. A) Bed B, Charnwood Forest, B) Bristy Cove/X-Ray surface, C) ‘D’ surface, D) H14/Johnson surface E) St. Shott’s surface, F) ‘E’ surface, G) Spaniard’s Bay surface. The univariate PCFs of H) Fractofusus and I) Charniodiscus from multiple surfaces are shown to demonstrate the similarity of their spatial distributions between localities. Where the best-fit model for the distribution represents a niche process (Table S3), it is drawn as a dashed line. Models indicating neutral processes are drawn as solid lines. Black line represents the random model. The grey area is the simulation envelope for 999 Monte Carlo simulations. The x-axis is the inter-point distance between organisms in metres. On the y-axis PCF=1 indicates complete spatial randomness (CSR), <1 indicates segregation, and >1 indicates aggregation. Different colors indicate different taxa as follows: Thectardis navy; Fractofusus light blue; Charnia bright yellow; Charniodiscus dark red; Aspidella light green; Bradgatia dark green; ‘Feather Duster’, light orange; Primocandelabrum dark orange; Trepassia dark purple; Beothukis bright pink; Pectinifrons dark blue; ‘Brushes’ brown; Avalofractus dark blue; Hylaecullulus light yellow.

**Figure 2.**
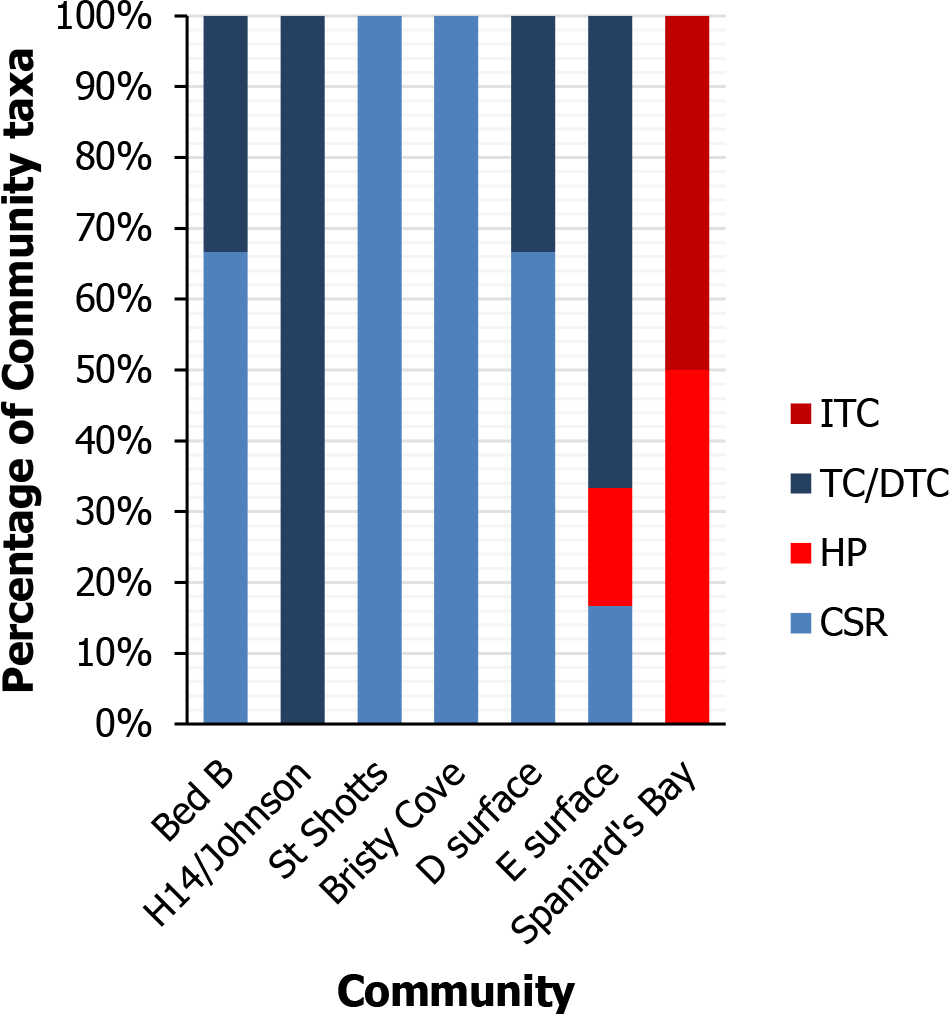
Proportion of best-fit univariate models by surface. The percentage of taxa with univariate spatial distributions that are best described by CSR, HP, TC (or DTC) and ITC models. CSR and TC are considered neutral models and shown in blue. HP and ITC are niche models, shown in red.

In order to gain some indication of whether taxa behave differently over large-spatial scales we compared the univariate spatial distributions of individual taxa at different sites to represent paleocommunities separated by large spatial and temporal scales. Four taxa are abundant on multiple bedding-planes (*Bradgatia*, *Charnia*, *Charniodiscus* and *Fractofusus*), and they all exhibit a consistent best-fit model (CSR, TC, TC and TC/DTC respectively) on each surface they are found on. Previous work has demonstrated that *Fractofusus* shows consistently the same spatial distributions across multiple surfaces (18) in different geological units (cf. H14 and ‘E’ surface, Fig. 1H). *Charniodiscus* also shows consistent spatial distributions (Fig. 1I), even when the communities were in temporally separated localities in different locations on Avalonia (e.g. *Charniodiscus* from the ‘E’ surface and Bed B). The consistency of these results suggests that the small-spatial-scale ecological behaviour of these taxa did not change over large spatial and temporal scales.

Two surfaces out of the five studied paleocommunities with more than one abundant taxon present exhibited only CSR bivariate distributions (St. Shott’s and the ‘D’ surface (22) Fig. 3, Table S3). The ‘E’ surface (22,44), Spaniard’s Bay and Bed B have exhibit non-CSR bivariate distributions (Fig. 3) indicating shared habitat associations. On Bed B the non-CSR bivariate distribution indicates shared habitat associations between the large *Charnia* and *Primocandelabrum* specimens (Fig. 3B, Table S3), as do the three non-CSR bivariate distributions on the ‘E’ surface (22, 44). For the *Primocandelabrum* ‘E’ surface and the Bed B non-CSR habitat associations, the large specimens had a segregated spatial distribution which corresponds to a reduced specimen density compared to CSR. This reduction in specimen density indicates that there were not enough resources to sustain all of the population and so is a niche process (44). On the ‘E’ surface, inter-specific segregations reduced specimen density by 25%, and aggregations increased specimen density by 56% (Fig. 3C, D). In contrast, intra-specific dispersal processes had a large effect on specimen density increasing their density between 250–600% (Fig. 1F). The two habitat associations on the ‘E’ surface (Feather Dusters–*Fractofusus* and Feather Dusters–*Charniodiscus*) were reflected in small bivariate aggregations (increase of 34% over distances under 0.2 m and 56% under 1.2 m respectively) with a reduction in their joint density at large spatial-scales (11% over 1 m and 13% over 2.1 m respectively; Fig. 3C, D). Similarly, habitat association between *Charnia* and *Primocandelabrum* on Bed B increased specimen density by 87%, whereas segregations reduced taxon density by 10%. Univariate dispersal-generated aggregations increased taxon density by 180–500% (Figs. 1A). The *Trepassia* – *Beothukis* bivariate distribution on Spaniard’s Bay is best modelled by the *Trepassia* best-fit model ITC model (Table S3) which is a Thomas Cluster model fitted onto the heterogeneous Poisson model background of *Beothukis* (Fig. 3A). This result demonstrates that there is a single habitat heterogeneity impacting both taxa on scales above 40 cm, but influencing *Beothukis* more strongly than *Trepassia* (Fig. 3G, although see *SI Appendix*). Across the paleocommunities, the bivariate habitat associations are much weaker in PCF magnitude than the univariate distributions (Figs 1 and 3), showing that the bivariate (niche) processes had less impact on spatial distributions than univariate (neutral) processes. These spatial distributions suggest that competition is rare, and that where it was present it was relatively weak in magnitude (Fig. 3; cf. refs. 22, 44).

**Figure. 3.**
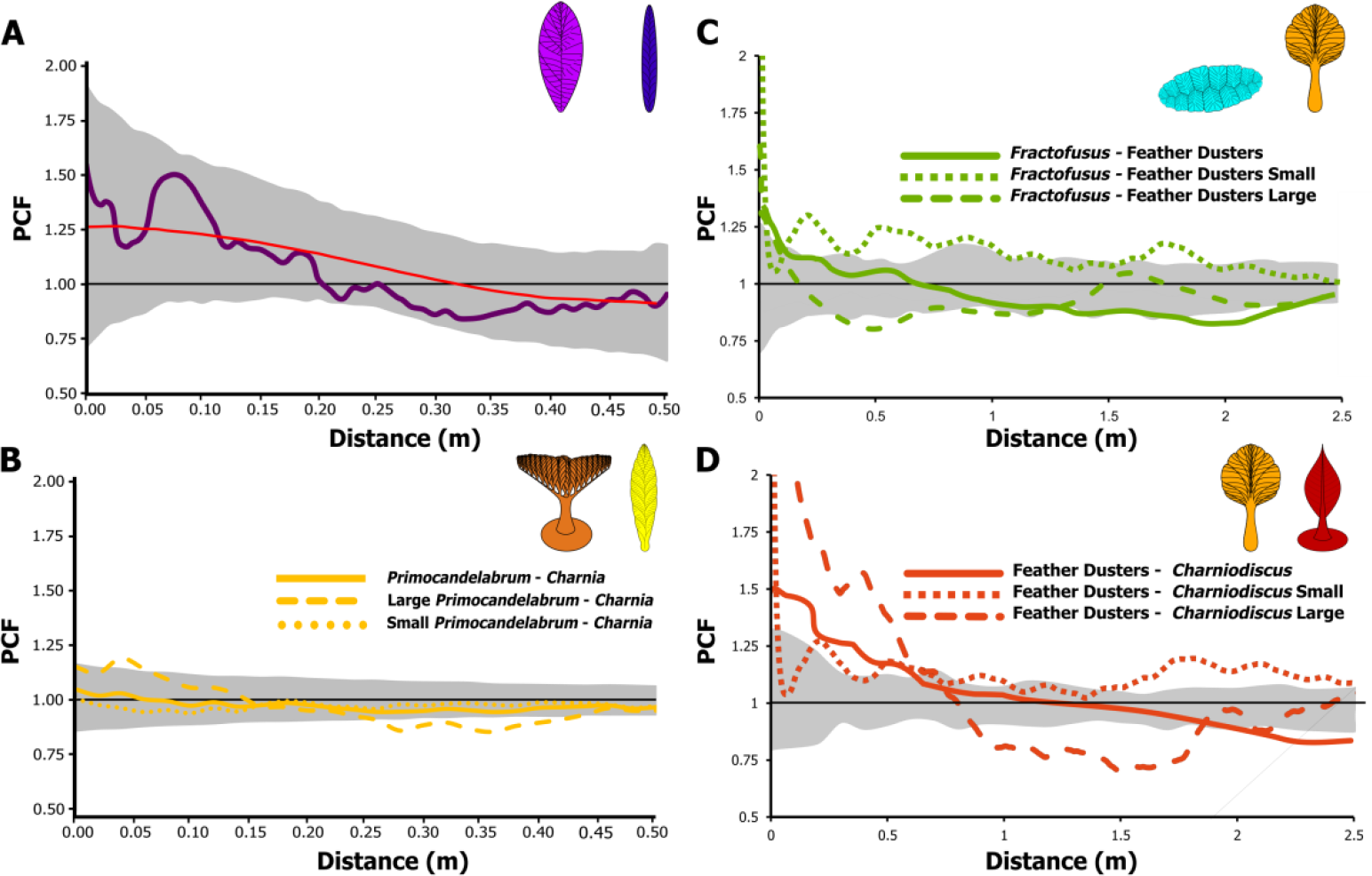
Bivariate PCF for the taxa which had non-random spatial distributions. The grey area is the simulation envelope for 999 Monte Carlo simulations. The x-axis is the inter-point distance between organisms in metres. On the y-axis, PCF=1 indicates complete spatial randomness (CSR) and is indicated by a black line, <1 indicates segregation, and >1 indicates aggregation. Red line is the modelled distribution. A) Trepassia and Beothukis from Spaniard’s Bay. The red line is the heterogeneous Poisson model. B) The non-CSR bivariate distribution of Charnia and Primocandelabrum from Bed B, Charnwood Forest shows randomly distributed small specimens (< 5.0 cm) and segregated large specimens(> 10.0 cm). C) Feather Dusters and Fractofusus and D) Feather Dusters and Charniodiscus from Mistaken Point ‘E’ Surface show aggregated small (< 3.0 cm) specimens and segregated large specimens (> 5.5cm). C and D reproduced from ref. 22.

## Discussion

Our results support combined theories of community assembly whereby niche and neutral theories are not mutually exclusive, but instead act along a continuum or spectrum, with differing extents of niche and neutral processes present in different circumstances (45, 46). The dominance of neutral best-fit univariate models, repetition of best-fit univariate models across different paleocommunities, and the rarity and weakness of bivariate niche best-fit models, all combine to provide strong evidence that neutral processes dominated Avalonian paleocommunities, with only limited niche-based influence. These neutral-dominated community dynamics contrast with those observed in the modern marine realm, where neutral processes are rare (47, 48).

The difference in dominance of niche versus neutral processes raises the question of whether the nature of community dynamics of Ediacaran metazoan-dominated paleocommunities was fundamentally different to those of the present day. The only other work on niche-neutral influences on paleocommunity assembly, is from the Quaternary (2.58 – 0.01 Ma), where fossil assemblages provides strong model and empirical support for environment-led (niche) models of assembly (49). There are some notable differences between Avalonian paleocommunities and extant marine communities. Avalonian paleocommunities appear to differ from the majority of extant marine systems in the extent of their ecological maturity, in that no more than three generations are seemingly preserved (22), though some paleocommunities include rare survivors (23) and/or evidence of secondary community succession (50). These characteristics suggest that the fossil communities are not always mature, many having been curtailed by high frequency incursions of sediment, limiting their maturity (22). Recent models show that community dynamics in small populations periodically subjected to disturbance events are dominated by neutral processes, implying a lack of small-spatial scale environmental control on their ecological dynamics (46).

The relative influence of niche versus neutral processes has been shown to be effected by the dispersal ranges of taxa within communities (52). Wide dispersal ranges increase the connectivity between populations, and so expand effective community size with the net effect of enhancing ecological selection (competition) and thereby increasing the relative importance of niche processes (52). The opposite is true when dispersal is limited, making these communities neutral dominated (52). Within the Ediacaran, the global distributions of some Avalonian taxa provide evidence that these taxa were capable of wide dispersal (51, 53). Further evidence of dispersal ranges is found in their spatial distributions (20, 44). For example, we can see in the PCF plots that *Fractofusus, Charniodiscus* and *Primocandelabrum* (Fig. 1A, H and I) have very short dispersal ranges, of <10 cm for *Fractofusus* and *Primocandelabrum* and <20 cm for *Charniodiscus*. However, *Fractofusus* was also capable of a waterborne propagule stage (20), and the global distribution of *Charniodiscus* suggests that it was as well, but that the waterborne phrase resulted in a minority of the population (20). Wide dispersal ranges are suggested by the global distribution of taxa such as *Charnia* (53) and also by the CSR and HP distributions of *Beothukis* and *Bradgatia* (Table S3). Six of the seven studied paleocommunities were dominated by taxa such as *Fractofusus* and *Charniodiscus* which predominantly exhibit limited local dispersal (Fig 1H, I; 18, 22, 44). The studied Ediacaran paleocommunities have comparatively small populations, experienced frequent disturbance events, and include many taxa with short dispersal ranges, so within this framework we would expect neutral processes to dominate. While the dominance of neutral-based processes within these paleocommunities differs significantly to the majority of the modern marine realm, the underlying dynamics are entirely consistent with models of assembly that include both niche and neutral processes, and are similar to those of modern communities subject to the same conditions. Thus, it is therefore likely that the fundamental mechanisms of metazoan community assembly were already in place in the Ediacaran Period, and so may have existed unchanged for ~570 million years.

In a similar manner to ecological processes, evolutionary processes can be categorised as niche (selection) or neutral (drift) processes (54). Selection (niche) processes are considered deterministic because external factors, such as limited resources, lead to competition in a predictable way: given a set of initial conditions, the organisms/communities will always respond to these conditions (environment) in the same way (54). By contrast, drift (neutral) processes are considered stochastic because they result from random fluctuations in population demography, so, given a set of the same initial conditions, different populations/communities may emerge. Hence, the observed dominance of neutral ecological processes in the Ediacaran Avalonian paleocommunities establishes that they are inherently stochastic/probabilistic with the possible implication that early metazoan diversification was not a systematic adaption to optimise survival under prevailing environmental conditions (which would be niche processes, and so deterministic). Instead, their existence under a stochastic regime would mean that diversification could have been merely driven by demographic differences resulting from random within-population. If this hypothesis is correct, and early metazoan evolution was stochastic, then this stochasticity may help to explain why neutral models of evolution can reproduce substantial macro-evolutionary trends such as the Cambrian Explosion (cf. 55), despite the known importance of niche processes in shaping evolution (e.g. 56).

## Conclusions

We have shown that paleocommunities of early macroscopic metazoans were overwhelming dominated by neutral ecological processes, with only limited and weak evidence for niche processes. Our results strongly contrast with modern marine systems, but because our Ediacaran paleocommunities have traits (short dispersal ranges, small populations and frequent disturbances) that are associated with neutral ecological models our results suggest that the fundamental mechanisms of community assembly may have been in place since the early stages of metazoan evolution. The dominance of neutral processes in these paleocommunities suggests that systematic adaptation of the Ediacaran organisms to their local environment may not have been the underlying driver of early metazoan diversification. Instead, it is possible that late Neoproterozoic metazoan diversification may result from stochastic demographic differences, with only limited environmental influence.

## Statement of Authorship

EGM conceived the project, designed the research, ran the analyses and wrote the first draft of the paper. All authors contributed to field data collection. SH developed the data post-processing protocol. SH and EGM processed field data. All authors were involved in writing the final manuscript.

## Data accessibility statement

Should the manuscript be accepted, the data supporting the results will be archived in Dryad, Figshare or Hal and the data DOI will be included at the end of the article.

## Acknowledgments

The Parks and Natural Areas Division (PNAD), Department of Environment and Conservation, Government of Newfoundland and Labrador provided permits to conduct research within the Mistaken Point Ecological Reserve (MPER) in 2010, 2016 and 2017. Readers are advised that access to MPER is by scientific research permit only. Contact PNAD for further information. Access to Bed B was kindly facilitated by Natural England and landowners in Charnwood Forest. This work has been supported by the Natural Environment Research Council [grant numbers NE/P002412/1 to EGM; NE/P002412/1 to CGK and PRW; and Independent Research Fellowship NE/L011409/2 to AGL], a Gibbs Travelling Fellowship from Newnham College, Cambridge and a Henslow Research Fellowship from Cambridge Philosophical Society to EGM. CGK also acknowledges a Research Studentship funded by the Cambridge Philosophical Society.

The authors declare no competing interests.

## Supplementary Information

**Figure S1.**
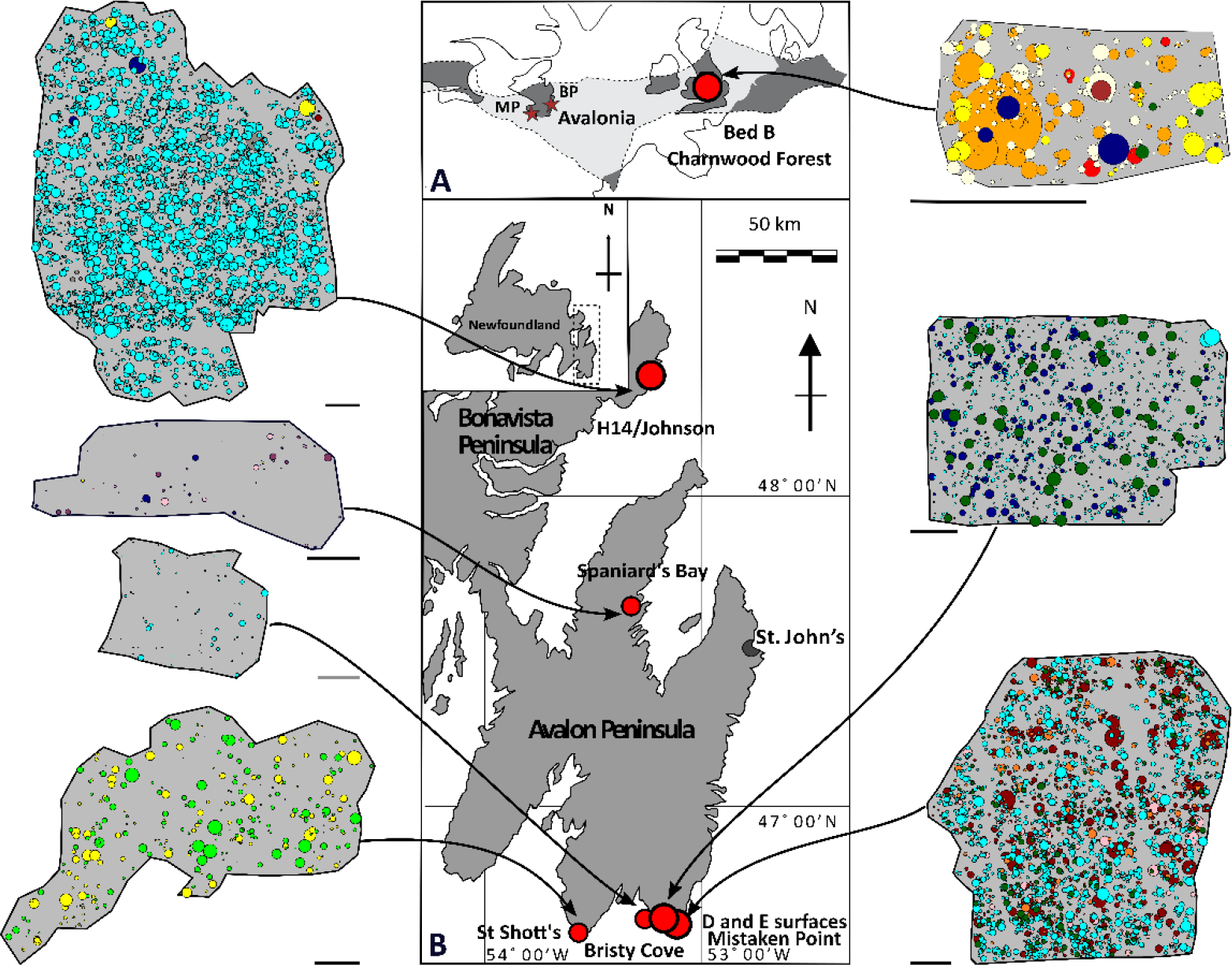
Locality map of study sites, showing: A, the relative location of sites with respect to the micro-continent of Avalonia (grey shading), highlighting the Mistaken Point Ecological Reserve (MP) and the Bonavista Peninsula (BP) in Newfoundland, and Bed B in Charnwood Forest, UK. B, the Newfoundland sites of the ‘D’, ‘E’, and Bristy Cove (X-Ray) surfaces, all in the Mistaken Point Ecological Reserve; the St. Shott’s (Sword Point) surface; and the H14/Johnson surface, Bonavista Peninsula (modified from ref. 22). Associated spatial maps for each locality show the positions of the fossil specimens, where the size of the circle indicates the vertical height. Black scale bar = 1 m, grey scale bar = 0.1 m. Different colors indicate different taxa as follows: Thectardis navy; Fractofusus light blue; Charnia bright yellow; Charniodiscus dark red; Aspidella light green; Bradgatia dark green; Feather Dusters light orange; Primocandelabrum dark orange; Trepassia dark purple; Beothukis bright pink; Pectinifrons dark blue; ‘Brushes’ brown; Avalofractus dark blue; Hylaecullulus light yellow.

**Table S1:** Summary data for each surface. Area is total area mapped. Total number of specimens includes non-abundant and taphomorph taxa. Density is of abundant taxa (>30 individuals) only. Minimum size consistently preserved is the modal height of small specimens, because specimens beneath this threshold exhibit discontinuous distributions (cf e.g. 79) so it is likely that specimens beneath this threshold may not have been preserved/mapped. *Note due to low specimen numbers on the Spaniard’s Bay surface, we included >15 specimens as abundant taxa (see SI appendix).

| Surface | Area mapped (m<sup>2</sup>) | Total number of specimens | Density (ind/m<sup>2</sup>) | Number of abundant (>30) taxa | Minimum size of consistently preserved features (cm) |
| --- | --- | --- | --- | --- | --- |
| *Bed B* | 115.16 | 761 | 6.61 | 3 | 1.20 |
| *MP E* | 82.44 | 2977 | 36.83 | 6 | 0.70 |
| *MP D* | 70.89 | 1402 | 22.29 | 3 | 1.20 |
| *H14* | 82.4 | 4235 | 52.55 | 1 | 1.50 |
| *BC* | 0.77 | 76 | 133.25 | 1 | 0.70 |
| *St. Shott's* | 50.93 | 480 | 7.20 | 2 | 1.20 |
| *Spaniard's Bay\** | 16.40 | 68 | 2.61 | 3 | 0.20 |

**Figure S2:**
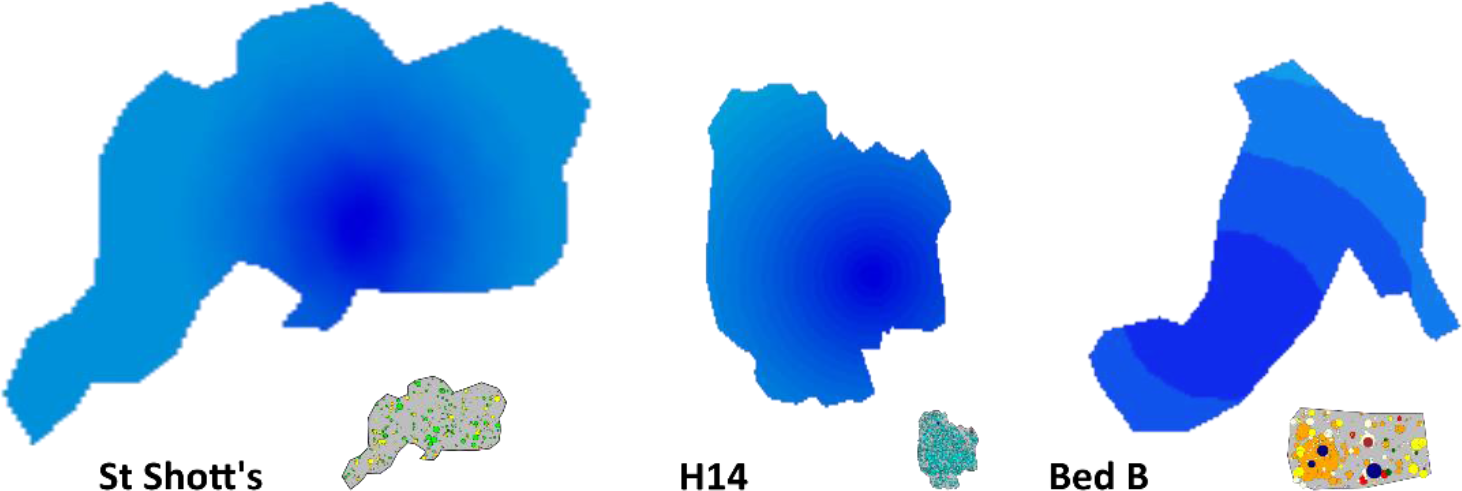
Model density maps of surfaces showing erosion biases. For each surface darker colors indicate higher modelled fossil density, and therefore lower presumed erosional rate, normalised for the density on each surface. Note that Bed B has a coarser pattern due to a relatively lower fossil density difference across the surface and that the full spatial map is not provided in full due to concerns about fossil theft.

**Table S2:** ΔAIC values for density models used to investigate erosional biases. *x* is parallel to strike, *y* parallel to dip, √*[(x−x_1_)^2^* + *(y−y_1_)^2^]* is the distance from the point of least erosion, and x_1_ and y_1_ are the co-ordinates of that point. These ΔAIC were used to determine the best-fit models. ΔAIC > 0 indicates that the model has a better fit to the data than completely spatially random model. Units are the centimetre co-ordinates of the spatial maps.

| Surface | X | y | $\sqrt{[(x-x_1)^2 + (y-y_1)^2]}$ | $x_1$ | $y_1$ |
| --- | --- | --- | --- | --- | --- |
| *Bed B* | 3.26 | **6.94** | 1.54 | 89 | 112 |
| *Bristy Cove* | -1.93 | -1.93 | -1.77 | 84 | 91 |
| *Mistaken Point D* | -3.37 | -12.29 | -12.51 | 52 | 939 |
| *Mistaken Point E* | -75.64 | -5.06 | -72.29 | 309 | 283 |
| *H14* | 43.26 | 37.69 | **87.34** | 98 | 70 |
| *St. Shott's* | -0.36 | 14.17 | **23.82** | 93 | 113 |
| *Spaniard's Bay* | 0.14 | -1.96 | -1.78 | 294 | 143 |

**Figure S3.**
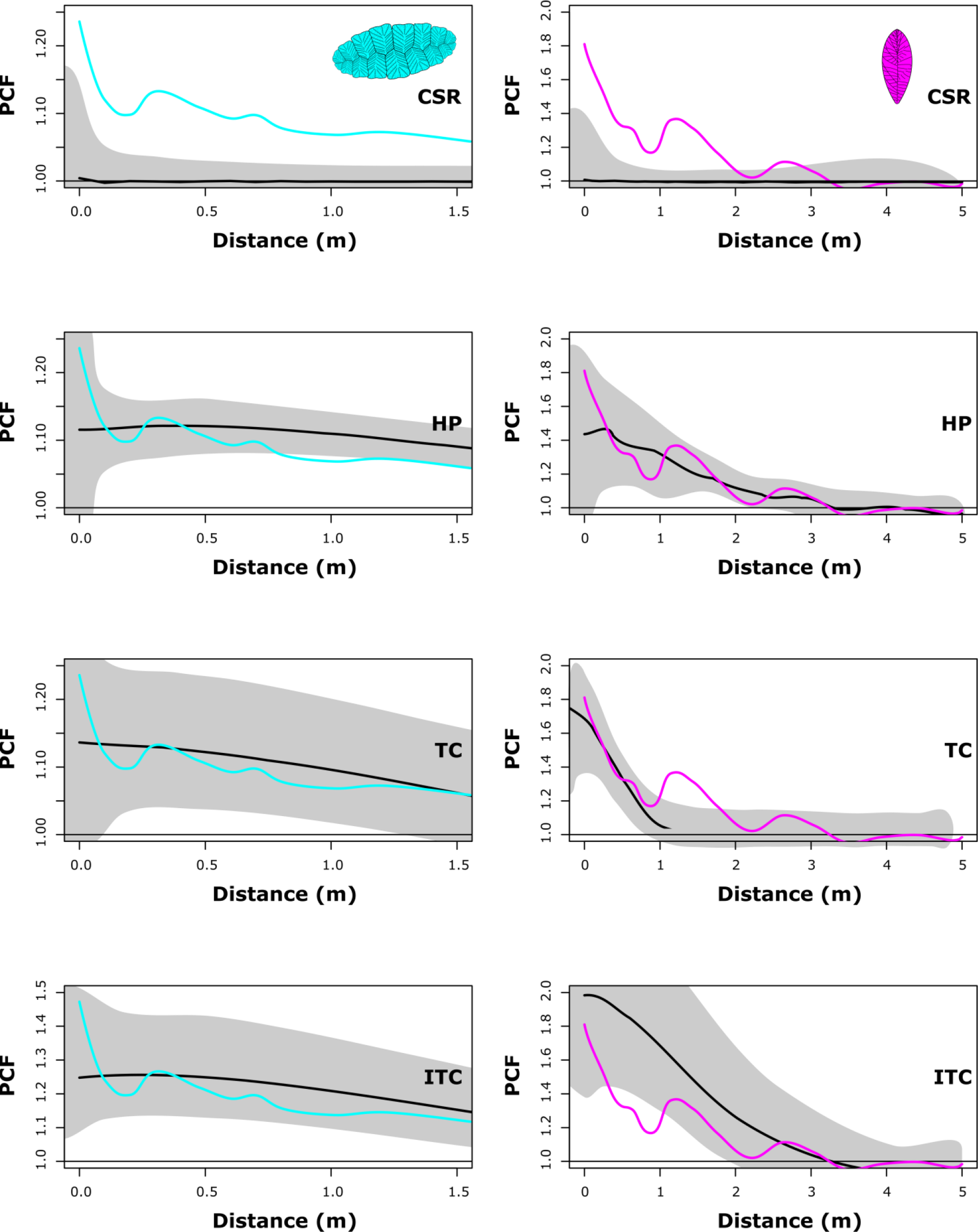
Univariate PCF analyses for Fractofusus on the ‘D’ surface (left) and Beothukis on the ‘E’ surface (right) under four different spatial models: CSR, HP, DTC, ITC (see text for model discussion). The model lines are black, the grey area represents the simulation envelope of 999 Monte Carlo simulations and the coloured lines are the observed spatial distributions For Fractofusus (left) the best-fit model is TC, whereas for Beothukis (right) it is HP because they follow the model best as evidenced by the Monte-Carlo simulations and p_d_ value (Table S3).

**Table S3:** Summary Table of Univariate PCF analyses. For the heterogeneous backgrounds, the moving window radius is 0.5 m, using the same taxon density as the taxon being modelled. *p*_*d*_ = 1 corresponds to a perfect fit of the model to the data, while *p*_*d*_ = 0 corresponds to no fit. Where observed data did not fall outside CSR Monte-Carlo simulation envelopes, no further analyses were performed, which is indicated by NA. σ: cluster radius, ρ: density of specimens, CSR: Complete spatial randomness, HP: Heterogeneous Poisson model, TC: Thomas Cluster model and ITC: inhomogeneous Thomas Cluster model. Note that if the cluster model is not a good fit, the *mean number in cluster* will not necessarily be appropriate.

| Surface | Taxon | Model fit | | | $P_d$ values | | | |
| --- | --- | --- | --- | --- | --- | --- | --- | --- |
| | | $\sigma$ | 100 $\rho$ | Mean number in cluster | CSR | HP | TC | ITC |
| **Bed B** | *Charniodiscus* | **2.906** | **0.7789** | **6** | 0.001 | 0.002 | 0.892 | 0.005 |
|  | *Primocandelabrum* | 21.47 | 0.1156 | 10 | 0.124 | 0.002 | 0.647 | 0.137 |
|  | *Charnia* | 12.027 | 0.4141 | 2 | 0.294 | 0.266 | 0.972 | 0.272 |
| **Bristy Cove** | *Fractofusus* | NA | NA | NA | 0.523 | NA | NA | NA |
| **Mistaken Point D** | *Fractofusus* | 4.312 | 0.638 | 3 | 0.001 | 0.51 | 0.77 | 0.21 |
|  | *Pectinifrons* | NA | NA | NA | 0.575 | NA | NA | NA |
|  | *Bradgatia* | NA | NA | NA | 0.664 | NA | NA | NA |
| **Mistaken Point E** | *Bradgatia* | NA | NA | NA | 0.44 | NA | NA | NA |
|  | *Charniodiscus* | 2.133 | 1.667 | 10 | 0.01 | 0.13 | 0.41 | 0.14 |
|  | *Beothukis* | 6.841 | 0.11 | 14 | 0.01 | 0.90 | 0.60 | 0.11 |
|  | Feather Dusters | 5.616 | 0.329 | 11 | 0.01 | 0.09 | 0.28 | 0.18 |
|  | *Thectardis* | 1.835 | 2.003 | 4 | 0.02 | 0.44 | 0.91 | 0.07 |
|  | *Fractofusus* | 11.842 | 0.39 | 25 | 0.01 | 0.03 | 0.32 | 0.35 |
| **H14** | *Fractofusus* | 7.906 | 0.192 | 2 | 0.001 | 0.003 | 0.760 | 0.004 |
| **St. Shott's** | *Aspidella* | 9.226 | 0.4032 | 4 | 0.011 | 0.016 | 0.963 | 0.015 |
|  | *Charnia* | 10.97 | 0.2443 | 2 | 0.257 | 0.225 | 0.331 | 0.271 |
| **Spaniard's Bay** | *Avalofractus* | 5.607 | 0.2151 | 2 | 0.087 | 0.142 | 0.665 | 0.307 |
|  | *Beothukis* | 36.70 | 0.834 | 8 | 0.401 | 0.810 | 0.530 | 0.308 |
|  | *Trepassia* | 7.148 | 0.1941 | 3 | 0.005 | 0.022 | 0.980 | 0.850 |

**Table S4.** Bivariate parameters for the best-fit models for the aggregated distributions. The parameters used for the shared source models (SS), linked cluster models (LCM) and linked double cluster models (LDCM). *p*_*d*_ = 1 corresponds to a perfect fit of the model to the data, while *p*_*d*_ = 0 corresponds to no fit at all.

| Surface | Taxon 1 | Taxon 2 | Model | Output values |  |  |
| --- | --- | --- | --- | --- | --- | --- |
| | | | | $\sigma$ (m) | Cluster Size | p-value |
| Spaniard's Bay | *Beothukis* | *Trepassia* | LCM | **36.70** | **8** | **0.797** |
|  | *Bradgatia* | *Beothukis* | SS | 7.43 | 14 | 0.166 |
| Bed B | *Charnia* | *Primocandelabrum* | LCM | **21.47** | **13** | **0.006** |
|  | *Charnia* | *Primocandelabrum* | SS | **8.571** | **17** | **0.786** |

## Supplementary Discussion

*Erosional analyses*

**Figure S4:**
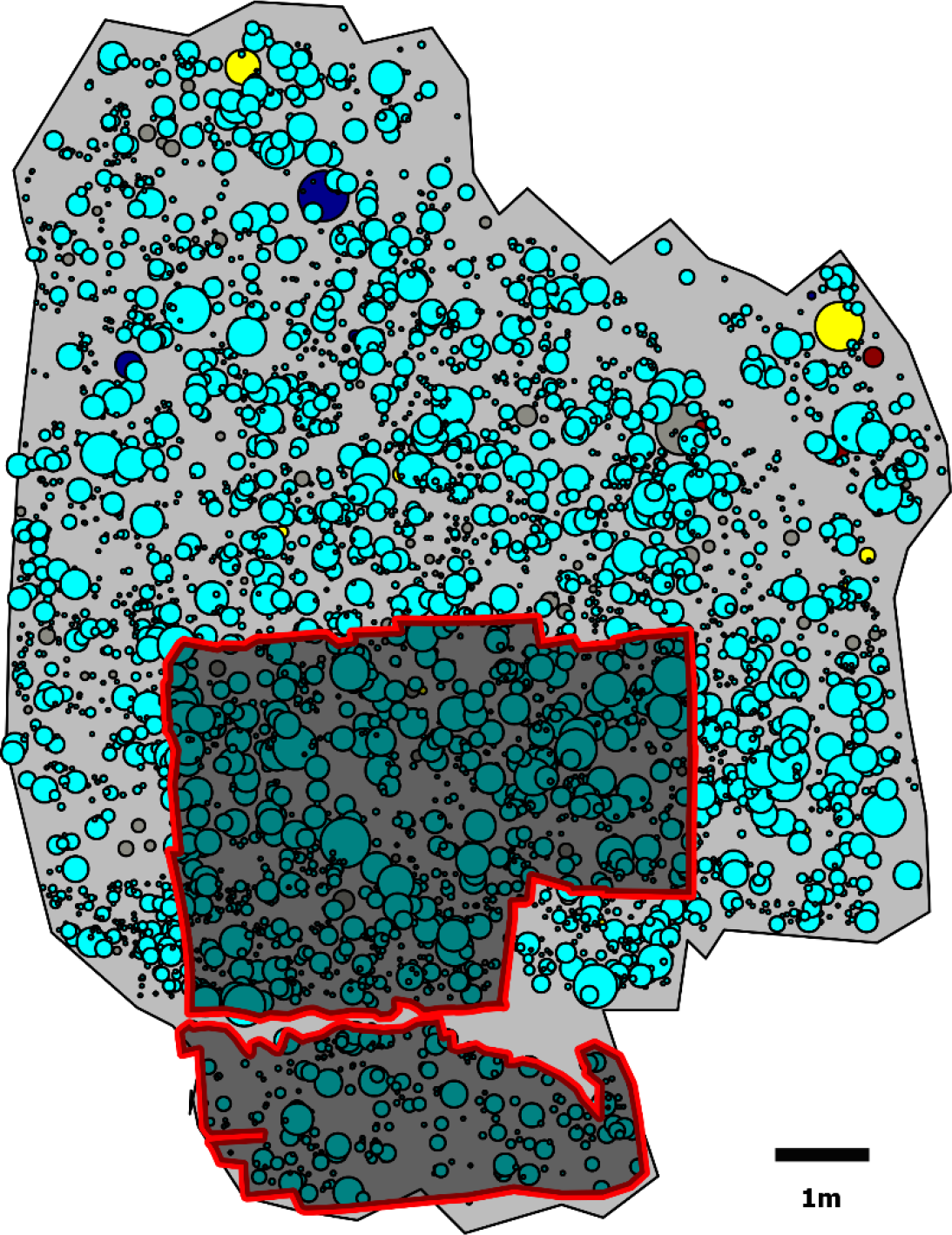
H14 spatial map showing the area mapped for Mitchell et al. 2015 study (red outline). Spatial maps show the positions of the fossil specimens, with the size of the circle indicated the length of the fossils (*Fractofusus*) or height (fronds) (indicated by a circle). Black scale bar = 1 m. Different colors indicate different taxa as follows: *Thectardis* navy; *Fractofusus* light blue; *Charnia* bright yellow; *Charniodiscus* dark red; Ivesheadiomorphs dark grey.

